# Autophagy deficiency in red pulp macrophages impairs their function and resistance to iron stress

**DOI:** 10.64898/2026.03.24.713972

**Authors:** Raquel Sal-Carro, Alice Lavanant, Marina Blanc, Gabriel Rojas-Jiménez, Blandine Maître, Christopher G. Mueller, Benjamin Voisin, Emmanuel L. Gautier, Frédéric Gros, Vincent Flacher

## Abstract

In mammals, most of the iron is found in the heme of red blood cells (RBCs), which must be recycled to support erythropoiesis in the bone marrow. Splenic red pulp macrophages (RPMs) play a crucial role in this process by phagocytosing senescent RBCs, metabolizing the heme and releasing iron back into the blood. Free cytoplasmic iron generates toxic reactive oxygen species, yet iron-specific adaptations of RPMs are not well documented. We previously reported that autophagy prevents ferroptosis in Langerhans cells, a cutaneous phagocyte subset. Thus, we hypothesized that autophagy may be important for the regulation of RPM metabolism and their maintenance of systemic iron homeostasis. To study this, we used *Atg5*^flox/flox^ and *Cd169*^cre^ mouse models to delete ATG5 in CD169^+^ macrophages, including RPMs. *Atg5*-deficient RPMs were decreased in number, and the remaining ones showed increased generation of toxic lipid peroxides. Spleens of *Atg5*^Δ*Cd169*^ mice were enlarged and contained more RBCs. Finally, autophagy impairment in RPMs exacerbated RBC loss in a model of phenylhydrazine-induced anemia. Our findings exemplify how dysregulation of macrophage metabolism alters their function and can disrupt tissue homeostasis upon challenge.

## INTRODUCTION

Splenic red pulp macrophages (RPMs) take up senescent or damaged red blood cells (RBCs) and metabolize hemoglobin to release iron into circulation^1^, thereby supporting erythropoiesis in the bone marrow^2^. Since RPMs have long lifespans and are maintained by local proliferation^3,4^, they are endowed with specific pathways to avoid iron-driven cytotoxicity^5^. Autophagy, a catabolic process where intracellular components are recycled by lysosomal degradation, may be part of this functional adaptation. Autophagy is especially relevant for long-lived cells as it maintains protein and organelle quality by limiting oxidative damage^6^. Moreover, autophagy is involved in iron metabolism by degrading ferritin to release iron in the cytoplasm^7,8^ and it limits ferroptosis in epidermal Langerhans cells^9^. Nonetheless, the role of autophagy in the homeostasis of RPMs is still poorly understood. Thus, we generated *Atg5*^Δ^*Cd169* mice, where autophagy is specifically disrupted in CD169-expressing macrophages, including RPMs.

## MATERIAL AND METHODS

### Mice

*Atg5*^Δ*Cd169*^ mice were generated by crossing *Atg5*^*flox*^ *mice*^10^ with the *Cd169*^*Cre*^ strain^11^, in which the *Cre* recombinase gene is inserted into the *Cd169* locus and is expressed under the control of its promoter. *Rubcn*^-/-^ mice were generated as previously described^9^. Genotyping is detailed in **Supplementary Material and methods**. All experiments were approved by the local ethics committee and carried out in conformity to national guidelines.

### Cell preparation and analysis

Spleens were digested for RPM analysis with 0.02mg/mL Collagenase D, 0.1mg/mL DNAse I and 0.8mg/mL Dispase II (Sigma) for 30min at 37ºC or crushed mechanically to analyze RBCs. To quantify spleen monocytes and neutrophils, the spleen was digested in 2,5mg/mL Collagenase D and 10 U/mL of DNAse I for 30min at 37ºC. Bone marrow cell suspensions were obtained from a femur that was flushed by centrifugation, as previously described^12^. For flow cytometry analysis, cells were stained with the antibodies listed in **Table S1**. To measure lipid peroxidation, cell suspensions were enriched using anti-CD11b magnetic bead separation (Miltenyi-Biotec) and incubated for 10min at 37ºC with 2mM Bodipy-C11 581/591 (Invitrogen) in PBS. Sample acquisition was performed on Gallios or CytoFlex LX cytometers (Beckman-Coulter). The number of RBCs and the hematocrit in blood and spleen were analysed by a Scil-Vet ABC automatic cell counter (Scil Animal Care Company).

### RNA extraction, cDNA synthesis and quantitative PCR

Splenocytes were enriched using biotinylated F4-80 **(Table S1)** and streptavidin magnetic nanobeads (Biolegend) according to the manufacturer’s instructions. After sorting spleen cells on an Aurora cell sorter (Cytek Biosciences), RNA was extracted using NucleoZOL (Macherey-Nagel). Retrotranscription and qPCR were performed using the RevertAid cDNA synthesis kit and SYBR-green master mix (ThermoFisher). Primers are listed in **Table S2**.

## RESULTS AND DISCUSSION

### RPMs of *Atg5*^Δ*Cd169*^ mice are in low numbers and undergo ferroptosis

To study the role of autophagy in the homeostasis of RPMs, we generated mice with a specific deletion of *Atg5* in CD169^+^ macrophages, including RPMs^13–15^. ATG5 is involved in autophagosome formation and its absence abrogates autophagy. We crossed *Cd169*^cre/+^ mice with *Atg5*^flox/flox^ mice to obtain control *Atg5*^WT^ or *Atg5*^Δ*Cd169*^ mice. Genotyping of the *Cd169* gene confirmed the insertion of the gene encoding Cre recombinase **(Figure S1A)** in *Atg5*^Δ*Cd169*^ mice. To confirm the invalidation of *Atg5* in RPMs, we sorted CD11b^low^ F4-80^high^ VCAM-1^+^ RPMs, as well as control myeloid cells (CD11b^high^ F4-80^+^ VCAM-1^-^) **(Figure S1B-C)**. *Spic*, an essential transcription factor for RPM differentiation and function^5,16^, was specifically expressed in RPMs but not in F4-80^+^ VCAM-1^-^ cells, indicating that only VCAM-1^+^ cells are *bona fide* RPMs. Whereas ATG5 seemed highly expressed in WT RPMs, RPMs of *Atg5*^Δ*Cd169*^ mice lacked *Atg5* expression, confirming efficient *Cd169*-dependent recombination in this subset **(Figure S1D-E)**.

We used flow cytometry to quantify RPMs in spleen cell suspensions **(Figure 1A)**. *Atg5*^Δ*Cd169*^ mice displayed a decrease of RPMs by half, both in percentage and in number per spleen **(Figure 1B)**. F4-80^+^ VCAM-1^-^ cells increased slightly in numbers but not in proportion **(Figure 1C)**. ATG5 is also implicated in cellular pathways other than autophagy, such as the LC3-associated phagocytosis (LAP), whereby LC3 is conjugated on the membrane of phagosomes to direct them towards lysosomes for degradation. Since LAP may be involved in the uptake of senescent or damaged RBCs by RPMs^17^, we sought to determine whether the phenotype observed in *Atg5*^Δ*Cd169*^ mice is autophagy-specific. *Rubcn*^-/-^ mice, which have impaired LAP but functional autophagy^18^, have the same levels of RPMs as their WT counterparts **(Figure S2A)**. Thus, defects of LAP do not account for the loss of *Atg5*-deficient RPMs.

**Figure 1.**
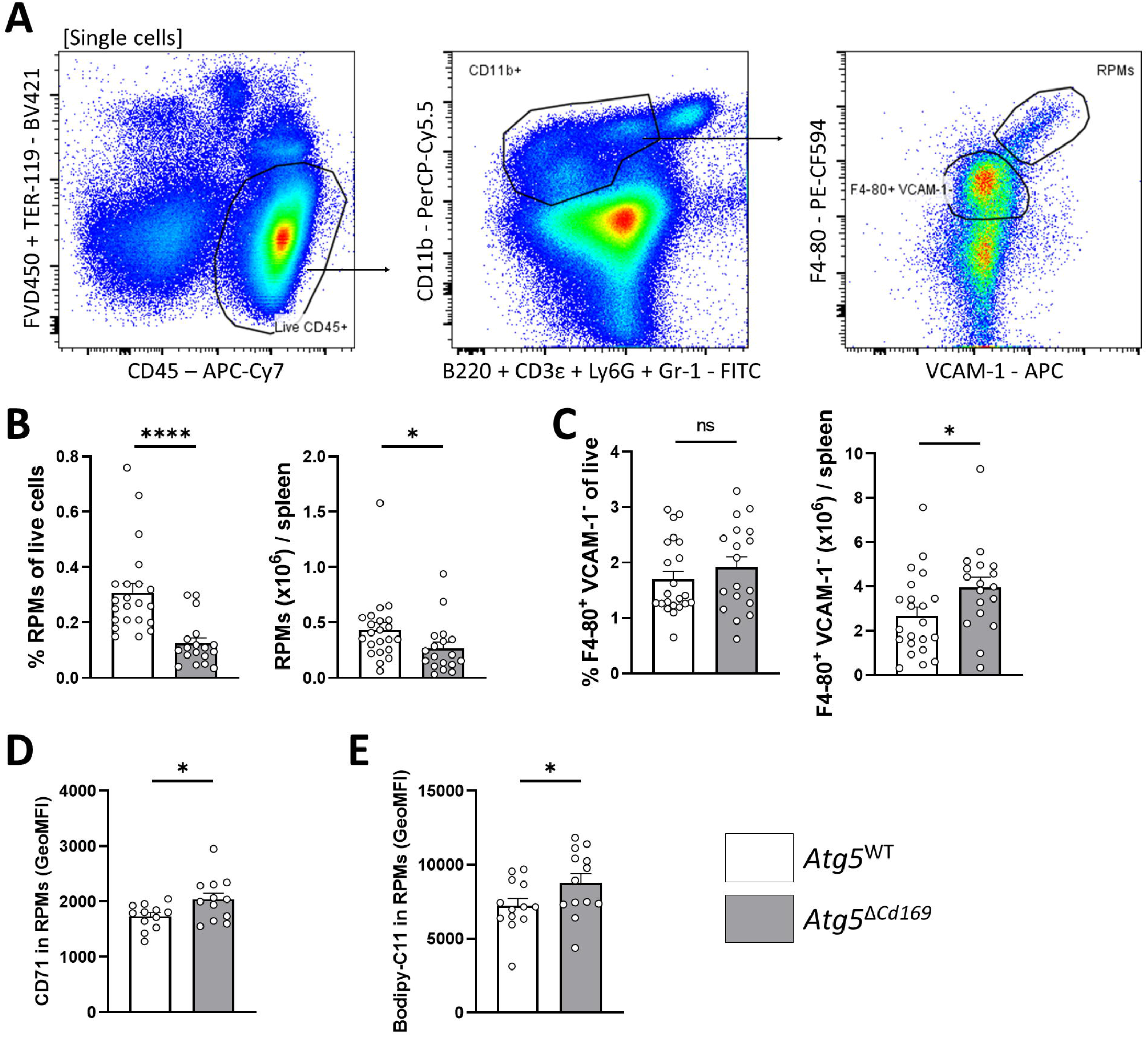
*Atg5*-deficient RPMs are decreased from the spleen and display signs of ferroptosis. **(A)** Strategy for gating RPMs and F4-80^+^ VCAM-1^-^ cells from spleen cell suspensions via flow cytometry. **(B)** Percentage of RPMs of live cells and number of RPMs per spleen obtained by flow cytometry analysis of splenic cell suspensions from *Atg5*^WT^ and *Atg5*^Δ*Cd169*^ mice (8-12 weeks old). **(C)** Percentage of F4-80^+^ VCAM-1^-^ cells and numbers per spleen obtained by flow cytometry analysis of spleen cell suspensions from *Atg5*^WT^ and *Atg5*^Δ*Cd169*^ mice. **(D)** Geometric mean fluorescence intensity (GeoMFI) of CD71 in RPMs analyzed by flow cytometry of RPM-enriched spleen cell suspensions. **(E)** GeoMFI of Bodipy-C11 staining in RPMs obtained by flow cytometry analysis of RPM-enriched spleen cell suspensions incubated with 2µM of Bodipy-C11 for 10min. Each point represents one mouse, error bars are SEM. Statistics: Mann-Whitney tests. ns, p>0.05; *, p<0.05; ****, p<0.0001.

RPMs recycle high quantities of iron from phagocytosed RBCs^2^. Iron accumulation catalyzes the oxidation of lipids, which can trigger cell death by ferroptosis^19^. In a mouse model of transfusion, excessive RBC phagocytosis leads to ferroptosis in RPMs^20^. We and others have reported that impaired autophagy is associated with such deleterious effects in other phagocytes^9,21^, which may explain the decrease in RPM numbers in *Atg5*^Δ*Cd169*^ mice. In line with this, surface expression of the iron transporter CD71 was increased in *Atg5*-deficient RPMs, and Bodipy-C11 assays confirmed higher levels of lipid peroxidation **(Figure 1D-E)**. Observing both of these hallmarks of ferroptotic cell death supports that autophagy is crucial to limit iron toxicity in RPMs.

### Spleens from *Atg5*^Δ*Cd169*^ mice are larger and accumulate RBCs

Spleen weight in *Atg5*^Δ*Cd169*^ mice was significantly increased as compared to *Atg5*^WT^ mice **(Figure 2A-B)**. As expected, the spleen weight of LAP-deficient *Rubcn*^-/-^ mice was the same as in control mice **(Figure S2B)**. Larger *Atg5*^Δ*Cd169*^ spleens were not a consequence of local inflammation, as they showed no detectable influx of neutrophils or monocytes **(Figure S3A-B)**. RPM-specific autophagy impairment did not cause systemic inflammation either **(Figure S3C-F)**. The total number of splenocytes was not higher in *Atg5*^Δ*Cd169*^mice **(Figure 2C)**. However, RBC numbers were higher in *Atg5*^Δ*Cd169*^ spleens **(Figure 2D)**, which could account for the increased spleen weight.

**Figure 2.**
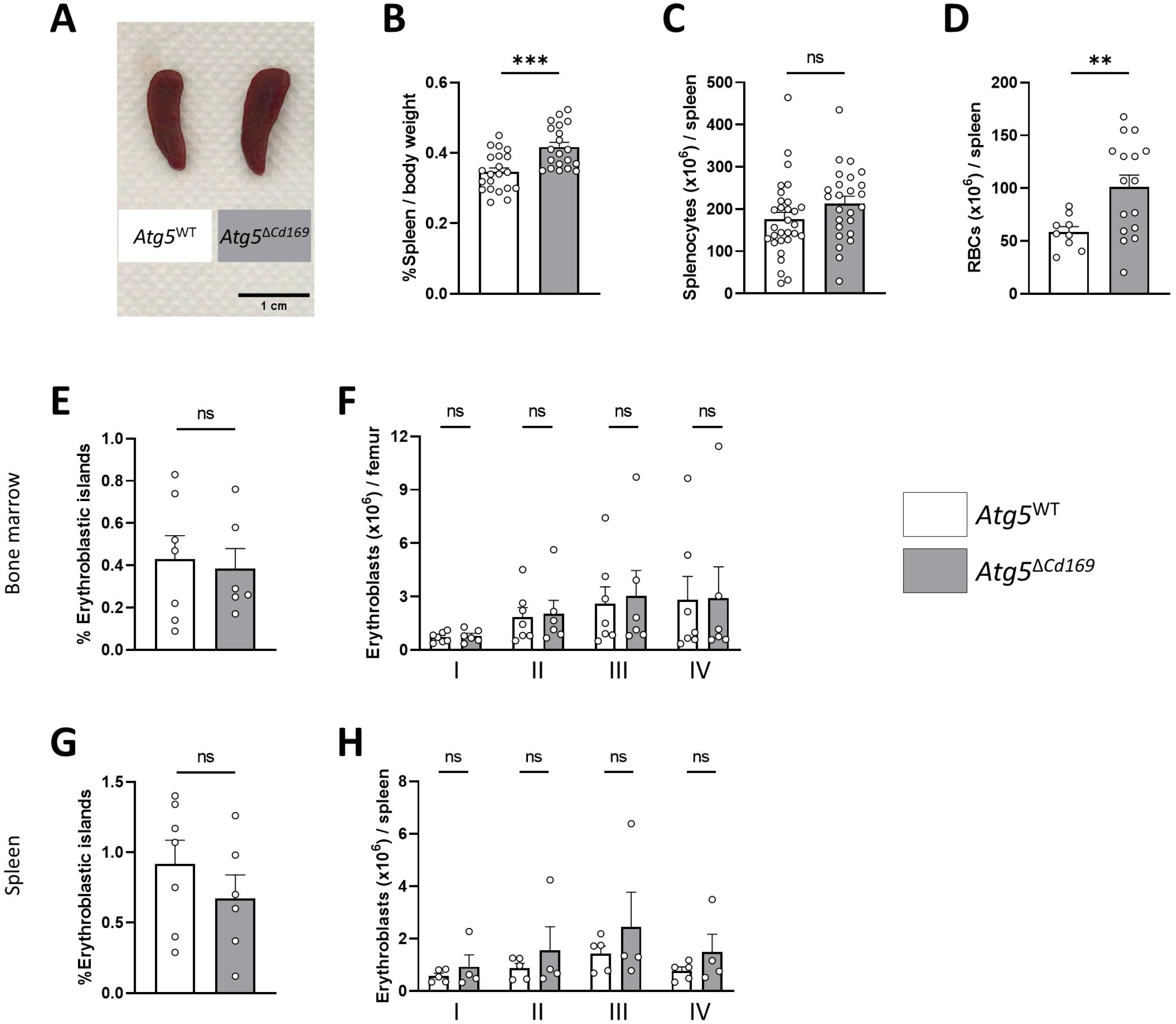
Spleens of *Atg5*^Δ*Cd169*^ mice are enlarged and accumulate erythrocytes. **(A)** Spleen pictures from 6-month-old *Atg5*^WT^ and *Atg5*^Δ*Cd169*^ mice. **(B)** Weight of the spleens relative to the weight of the mice (8-12 weeks old). **(C)** Number of splenocytes per spleen, counted in spleen cell suspensions after the lysis of RBCs. **(D)** Numbers of RBC per spleen, obtained by analysis of mechanically disrupted spleens. **(E)** Percentage of F4-80^+^ TER-119^+^ erythroblastic islands determined by flow cytometry in femur bone marrow cell suspensions. **(F)** Quantification of four types of erythroblastic progenitors per femur in bone marrow cell suspensions (I, Proerythroblasts; II, Basophilic erythroblasts; III, Polychromatic erythroblasts; IV, Orthochromatic erythroblasts). **(G)** Percentage of erythroblastic islands determined by flow cytometry in spleen cells suspensions. **(H)** Quantification of four types of erythroblastic progenitors per spleen weight. Each point represents one mouse, error bars are SEM. Statistics: Mann-Whitney test for B-E and G; mixed effects analysis and Sidak’s multiple comparison test for F and H. ns, p>0.05; **, p<0.01; ***, p<0.001.

Considering their role in RBC homeostasis, the reduction of RPMs might cause senescent or damaged RBCs to accumulate in the blood or in the spleen. Yet, in *Atg5*^Δ*Cd169*^ mice, the reduction of RPMs did not change RBC numbers in the blood **(Figure S4A-B)**. Phosphatidylserine, a hallmark of eryptosis, and autologous IgG molecules, which opsonize damaged RBCs^22^, are involved in the detection of senescent RBCs. *Atg5*^Δ*Cd169*^ percentages of eryptotic, dead and opsonized RBCs in the blood **(Figure S4C-H)** and the spleen **(Figure S4I-N)** were similar to *Atg5*^WT^ mice. Thus, increased RBCs in the spleen after RPM reduction does not translate into alterations of blood RBCs.

Decreased erythrophagocytosis may promote erythropoietic activity, leading to splenomegaly^14^. Erythroblastic islands contain TER-119^+^ RBCs associated with F4-80^+^ macrophages, which also express CD169^13,14^ **(Figure S5A, C)**, and erythroblasts at different stages of differentiation **(Figure S5B, D)**. The quantities of erythroblastic islands or the numbers of any of the erythroblast subsets were unchanged in the bone marrow **(Figure 2E-F)**. Similarly, there was no sign of extramedullary hematopoiesis in the spleen **(Figure 2G-H)**.

Overall, despite RPM reduction and splenomegaly, the numbers of circulating RBCs remained unaltered in *Atg5*^Δ*Cd169*^ mice, and RBCs did not appear damaged or senescent. Thus, the remaining RPMs might suffice to maintain erythrocyte homeostasis in the steady state. Still, the splenomegaly suggested a lower rate of clearance by ATG5-deficient RPMs, which could become deleterious in a pathological context.

### *Atg5* deletion in RPMs exacerbates the loss of RBCs upon phenylhydrazine-induced anemia

We hypothesized that the loss of RPMs may have consequences in hemolytic anemia, when many RBCs are simultaneously damaged and must be quickly removed from circulation. Acute anemia can be induced by treatment with phenylhydrazine (PHZ)^14^ **(Figure 3A)**. PHZ-treated mice lost weight until day 4, and fully recovered by day 7 **(Figure 3B)**. PHZ caused RBC lysis, with the lowest hematocrit and RBC levels reached 3 days after the injection **(Figure 3C-D)**.

**Figure 3.**
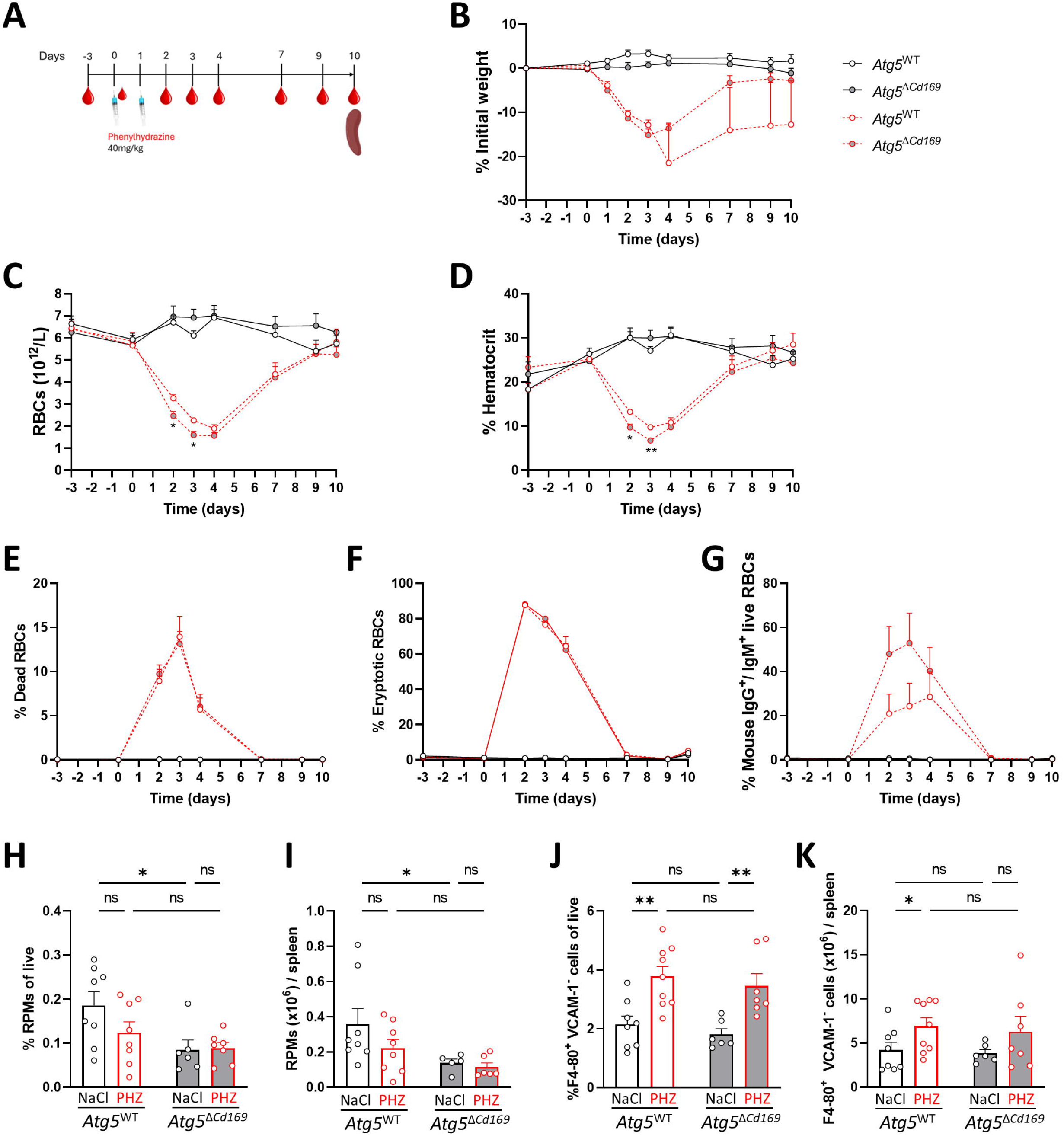
*Atg5* deletion in RPMs exacerbates the loss of RBCs upon phenylhydrazine-induced anemia. **(A)** Schematic representation of phenylhydrazine (PHZ) treatment and collection of blood and spleen. **(B)** Percentage of the initial weight, determined 3 days before PHZ treatment, measured until 10 days after the first PHZ injection for *Atg5*^WT^ and *Atg5*^Δ*Cd169*^ mice. **(C)** Number of RBCs and **(D)** hematocrit in the blood of *Atg5*^WT^ and *Atg5*^Δ*Cd169*^ mice injected with PHZ or NaCl (control) over the course of the experiment. **(E-G)** Percentage of dead (Annexin-V^+^ FVD450^+^) **(E)**, eryptotic (Annexin-V^+^ FVD450^-^) **(F)** and opsonized (mouse IgG^+^/IgM^+^ live RBCs) **(G)** RBCs in the blood of *Atg5*^WT^ and *Atg5*^Δ*Cd169*^ mice injected with PHZ or NaCl and analyzed by flow cytometry. **(H)** Percentage of RPMs out of live cells. **(I)** Number of RPMs per spleen in *Atg5*^WT^ and *Atg5*^Δ*Cd169*^ mice 10 days after the first injection of PHZ or NaCl. **(J)** Percentage of F4-80^+^ VCAM-1^-^ cells out of live cells. **(K)** Number of F4-80^+^ VCAM-1^-^ cells per spleen in *Atg5*^WT^ and *Atg5*^Δ*Cd169*^ mice 10 days after the first injection of PHZ or NaCl. Each point in B-F represents the average of 6-12 mice, error bars are the SEM. Each point in H-I represents an individual mouse, error bars are SEM. Statistics: mixed-effects analysis (REML) and Tukey’s multiple comparison test for B-F; 2-way ANOVA and Tukey’s multiple comparison test for H-I. ns, p>0.05; *, p<0.05; **, p<0.01. The asterisks indicated in B-G represent the p-value of the comparison between treated *Atg5*^WT^ and *Atg5*^Δ*Cd169*^ mice, non-significant differences were omitted.

In *Atg5*^Δ*Cd169*^ mice, RBC loss was exacerbated 2 and 3 days after injection, as shown by significantly lower levels of hematocrit and RBCs. However, their recovery was similar to WT mice. Concomitant with RBC loss, the proportion of dead, eryptotic and IgG/IgM-bound RBCs sharply increased and peaked at day 3 **(Figure 3E-G)**. Yet, no differences between *Atg5*^Δ*Cd169*^ and *Atg5*^WT^ mice were observed in the percentage of dead or eryptotic RBCs **(Figure 3E-F)**, while opsonized RBCs only showed a non-significant increase in *Atg5*^Δ*Cd169*^mice at days 2 and 3 **(Figure 3G)**. Finally, in spleens monitored after recovery (day 10), PHZ did not cause significant changes in RPMs **(Figure 3H-I)**. Conversely, F4-80^+^ VCAM-1^-^ cells increased upon PHZ injection for both *Atg5*^WT^ and *Atg5*^Δ*Cd169*^ mice **(Figure 3J-K)**. This population could correspond to the previously described CD11b^high^ F4-80^+^ VCAM-1^-^ subset, which differentiates into RPMs upon exposure to heme^5^.

Altogether, *Atg5* deficiency in RPMs results in their partial depletion and functional impairment, temporarily aggravating the loss of RBCs in PHZ-induced anemia. Adding to a growing body of evidence in different tissue-resident macrophages^9,21,23^, we conclude that the survival and function of RPM require autophagy, which should be further examined in pathological conditions^24^.

## ACKNOWLEDGEMENTS

VF is employed by the Centre National de la Recherche Scientifique and obtained funding from the Agence Nationale de la Recherche (ANR/ERAPerMed BATMAN (ANR-18-PERM-0001), AUTOMATE (ANR-20-CE15-0018-01, with partners FG and ELG), LabCom INCREASE (ANR-22-LCV2-0009-01)) and the Institut national du cancer (INCA EN-HOPE SMART4CBT). BV is employed by University of Strasbourg and is recipient of ANR JCJC DEEPENS. RSC was supported by ANR AUTOMATE and DEEPENS. GRJ was supported by Marie Skłodowska-Curie Action EURIdoc (EU-MSCA-COFUND-DP N°101034170). Experimental schemes were created with Biorender.com.

## AUTHORSHIP CONTRIBUTIONS

RSC, FG and VF conceptualized the research. RSC, AL, MB performed experiments and analyzed the data. Additional analyses of circulating blood cells were done by GRJ and BM. RSC and VF wrote the manuscript, with critical contributions by ELG, FG, BV and CM.

## SUPPLEMENTARY FIGURE LEGENDS

**Figure S1. *Atg5* is specifically deleted in RPMs in *Atg5***^Δ***Cd169***^ **mice. (A)** PCR of *Atg5*^WT^ (*Cd169*^*+/+*^ *Atg5*^*flox/flox*^) and *Atg5*^Δ*Cd169*^ (*Cd169*^*Cre/+*^ *Atg5*^*flox/flox*^) mice to detect the fragment amplified from the CRE recombinase gene (766bp, compared to the 519bp fragment in WT mice). **(B)** Gating strategy used to sort RPMs and F4-80^+^ VCAM-1^-^ cells from F4-80-enriched spleen cell suspensions. **(C)** Backgating of RPMs and F4-80^+^ VCAM-1^-^ populations onto CD11b^+^ lineage^-^ spleen cells. **(D-E)** qPCR to detect the expression of *Spic* **(D)** and *Atg5* **(E)** relative to *Actb* from sorted F4-80^+^ VCAM-1^-^ cells and RPMs. N=2; n=3. Each point represents one mouse. Statistics: Two-way ANOVA and Tukey’s multiple comparisons test. ns, p>0.05; *, p<0.05; **, p<0.01.

**Figure S2. RPMs are not impacted by the loss of *Rubcn* expression. (A)** Percentage of RPMs of live cells in spleen cell suspensions analyzed by flow cytometry. **(B)** Spleen weight relative to mouse weight of *Rubcn*^+/+^, *Rubcn*^+/-^ and *Rubcn*^-/-^ mice. Each point represents one mouse. Error bars are SEM. Statistics: Kruskal-Wallis one-way ANOVA and Dunn’s multiple comparison test. ns, p>0,05, N=1.

**Figure S3. The reduction of RPMs does not cause local or systemic inflammation. (A)** Number of neutrophils (CD45^+^ VCAM-1^-^ CD11b^+^ Ly6G^+^) in spleen cell suspensions per gram of digested spleen obtained by flow cytometry analysis. **(B)** Number of monocytes (CD45^+^ VCAM-1^-^ CD11b^+^ Ly6C^+^) in spleen cell suspensions per gram of digested spleen. **(C-F)** Quantification of white blood cells **(C)**, lymphocytes **(D)**, neutrophils **(E)** and monocytes **(F)**. Each point represents one mouse. Error bars are SEM. Statistics: Mann-Whitney test, ns, p>0.05.

**Figure S4. RPM decrease does not affect RBC homeostasis in the steady state. (A-B)** Quantification of RBC numbers **(A)** and hematocrit levels **(B)** in the blood of *Atg5*^WT^ and *Atg5*^Δ*Cd169*^ mice. **(C)** Representative flow cytometry staining of Annexin-V and viability dye (FVD450) in blood RBCs. **(D)** Percentage of dead (FVD450^+^ Annexin-V^+^) RBCs in the blood of *Atg5*^WT^ and *Atg5*^Δ*Cd169*^ mice. **(E)** Percentage of eryptotic (FVD450^-^ Annexin-V^+^) RBCs in the blood of *Atg5*^WT^ and *Atg5*^Δ*Cd169*^ mice. **(F)** Representative flow cytometry staining of surface-bound autologous IgG and viability dye (FVD450) in blood RBCs. **(G)** Percentage of IgG-bound dead (FVD450^+^ Mouse IgG^+^) RBCs in the blood of *Atg5*^WT^ and *Atg5*^Δ*Cd169*^mice. **(H)** Percentage of live IgG-bound (FVD450^-^ Annexin-V^+^) RBCs in the blood of *Atg5*^WT^ and *Atg5*^Δ*Cd169*^ mice. **(I)** Representative flow cytometry staining of Annexin-V and FVD450 in RBCs obtained from mechanically disrupted spleen cell suspensions. **(J)** Percentage of dead (FVD450^+^ Annexin-V^+^) RBCs in the spleen of *Atg5*^WT^ and *Atg5*^Δ*Cd169*^ mice. **(K)** Percentage of eryptotic (FVD450^-^ Annexin-V^+^) RBCs in the spleen of *Atg5*^WT^ and *Atg5*^Δ*Cd169*^ mice. **(L)** Representative flow cytometry staining of surface-bound autologous IgG and viability dye (FVD450) in spleen RBCs. **(M)** Percentage of IgG-bound dead (FVD450^+^ Mouse IgG^+^) RBCs in the spleen of *Atg5*^WT^ and *Atg5*^Δ*Cd169*^mice. **(N)** Percentage of live IgG-bound (FVD450^-^ Annexin-V^+^) RBCs in the spleen of *Atg5*^WT^ and *Atg5*^Δ*Cd169*^ mice. Each point represents one mouse. Error bars are SEM. Statistics: Mann-Whitney test. ns, p>0,05.

**Figure S5. Gating strategy for the identification of erythroblastic islands and progenitors in spleen and bone marrow. (A)** Representative flow cytometry staining of erythroblastic islands (F4-80^+^ TER-119^+^ multiplets) in bone marrow cells. **(B)** Representative flow cytometry staining of erythroblastic progenitors (CD11b^-^ CD45^-^) in the bone marrow (I, Proeythroblasts; II, Basophilic erythroblasts; III, Polychromatic erythroblasts; IV, Orthochromatic erythroblasts).

(C)Representative flow cytometry staining of erythroblastic islands in spleen cell suspensions.

(D)Representative flow cytometry staining of erythroblastic progenitors in the spleen.

## SUPPLEMENTARY MATERIAL AND METHODS

**Table S1.** Antibodies used for flow cytometry.

| Target | Clone | Conjugate | Reference | Supplier |
| --- | --- | --- | --- | --- |
| Annexin-V | N/A | FITC | 556419 | BD |
| CD71 | C2 | FITC | 553266 | BD |
| TER-119 | TER-119 | AF488 | 116215 | Biolegend |
| B220 | RA3-6B2 | FITC | 103206 | Biolegend |
| CD3ε | 145-2C11 | FITC | 553062 | BD |
| GR-1 | RB6-8C5 | FITC | 108406 | Biolegend |
| Ly6G | 1A8 | FITC | 551460 | BD |
| NK1.1 | PK136 | AF488 | 108718 | Biolegend |
| F4-80 | BM8 | PE | 123110 | Biolegend |
| CD45 | 30F11 | PE-Cy7 | 103114 | Biolegend |
| F4-80 | BM8 | Biotin | 13-4801-82 | eBioscience |
| CD11b | M1/70 | PerCP-Cy5.5 | 101228 | Biolegend |
| CD11c | HL3 | PE-Cy7 | 558079 | BD |
| VCAM-1 | 429 | APC | 105718 | Biolegend |
| CD45 | 30-F11 | APC-Cy7 | 103116 | Biolegend |
| TER-119 | TER-119 | BV421 | 116234 | Biolegend |
| B220 | RA3-6B2 | BV421 | 103239 | Biolegend |
| CD3ε | 145-2C11 | BV421 | 100335 | Biolegend |
| GR-1 | RB6-8C5 | BV421 | 108434 | Biolegend |
| CD45 | 30F11 | BV785 | 103149 | Biolegend |
| CD47 | miap301 | PE-Dazzle 594 | 127521 | Biolegend |
| Mouse IgG | Polyclonal | AF647 | A-21235 | ThermoFisher |
| Mouse IgM | RMM-1 | AF647 | 406525 | Biolegend |
| F4-80 | BM8 | BV650 | 123149 | Biolegend |
| MHC-II | M5/114.15.2 | AF700 | 107622 | Biolegend |
| B220 | RA3-6B2 | BUV737 | 612839 | BD |
| CD3ε | 145-2C11 | BUV737 | 612771 | BD |
| CD44 | IM7 | PE-Cy7 | 560569 | BD |
| DAPI | N/A | N/A | D3571 | Invitrogen |
| Fixable Viability dye eFluor 450 (FVD450) | N/A | N/A | 65-0863-14 | Invitrogen |
| Fixable Viability dye eFluor 780 (FVD780) | N/A | N/A | 65-0865-14 | Invitrogen |
| ViaKrome808 Fixable Viability Dye | N/A | N/A | C36628 | Beckman Coulter |
| Streptavidin | N/A | PE-CF594 | 562284 | BD |

**Table S2.**
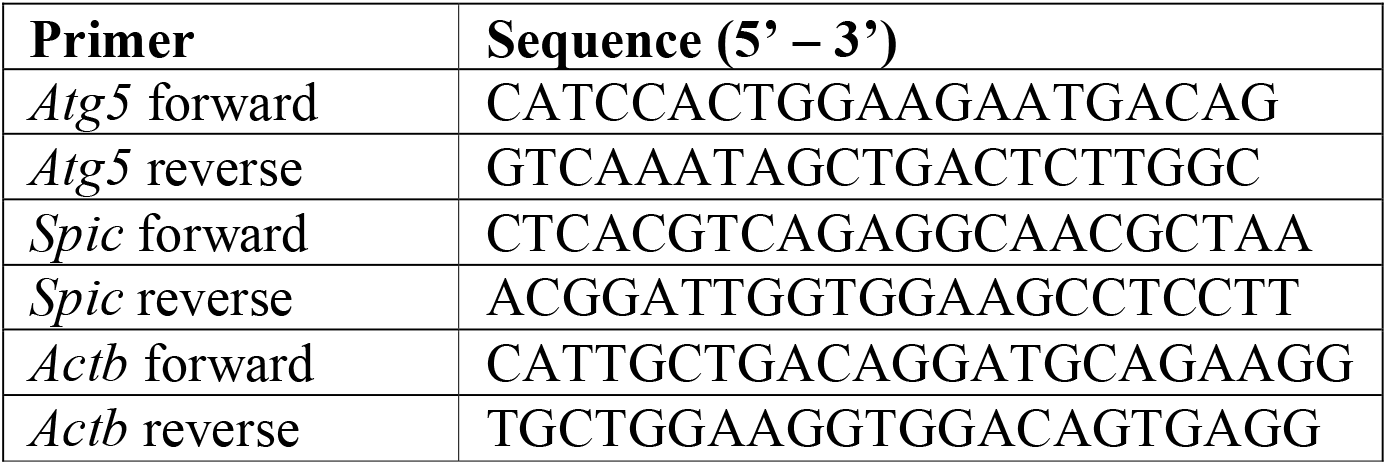
Primers for qPCR.

### Mice genotyping

Mice were genotyped at 5-11 days old for *Cd169* and *Atg5* alleles. The DNA was extracted from the tip of the tail with the kit Extract-N-Amp (Sigma Aldrich) according to the manufacturer’s instructions. PCR was performed using MyTaqTM HS Red Mix (Ozyme) using the following primers: *Atg5* exon 3: 5’-GAATATGAAGGCACACCCCTGAAATG-3’; *Atg5* CHECK: 5’-ACAACGTCGAGCACAGCTGCGCAAGG-3’; *Atg5* SHORT: 5’-GTACTGATAATGGTTTAACTCTTGC-3’; *Cd169* forward: 5’-GCTTACGGTGCTTGCTGGAT-3’; *Cd169* reverse: 5’-CATAGTCTAGGCTTCTGTGC-3’; and *Cd169*-cre reverse: 5’-AGGGACACAGCATTGGAGTC-3’. The PCR program started with 4min at 95ºC, followed by 34 cycles of 30s at 94ºC, 30s at 62ºC and 1min at 72ºC and a final incubation of 7min at 72ºC. The amplified fragments were analyzed by electrophoresis in a 1.5% agarose gel.

## REFERENCES

1. Kovtunovych, G., Eckhaus, M. A., Ghosh, M. C., Ollivierre-Wilson, H. & Rouault, T. A. Dysfunction of the heme recycling system in heme oxygenase 1-deficient mice: effects on macrophage viability and tissue iron distribution. Blood 116, 6054–6062 (2010).

2. Klei, T. R. L., Meinderts, S. M., van den Berg, T. K. & van Bruggen, R. From the Cradle to the Grave: The Role of Macrophages in Erythropoiesis and Erythrophagocytosis. Front Immunol 8, 73 (2017).

3. Hashimoto, D. et al. Tissue-Resident Macrophages Self-Maintain Locally throughout Adult Life with Minimal Contribution from Circulating Monocytes. Immunity 38, 792–804 (2013).

4. Yona, S. et al. Fate Mapping Reveals Origins and Dynamics of Monocytes and Tissue Macrophages under Homeostasis. Immunity 38, 79–91 (2013).

5. Haldar, M. et al. Heme-Mediated SPI-C Induction Promotes Monocyte Differentiation into Iron-Recycling Macrophages. Cell 156, 1223–1234 (2014).

6. Gómez-Virgilio, L. et al. Autophagy: A Key Regulator of Homeostasis and Disease: An Overview of Molecular Mechanisms and Modulators. Cells 11, 2262 (2022).

7. Bellelli, R. et al. NCOA4 Deficiency Impairs Systemic Iron Homeostasis. Cell Reports 14, 411–421 (2016).

8. Mancias, J. D., Wang, X., Gygi, S. P., Harper, J. W. & Kimmelman, A. C. Quantitative proteomics identifies NCOA4 as the cargo receptor mediating ferritinophagy. Nature 509, 105–109 (2014).

9. Arbogast, F. et al. Epidermal maintenance of Langerhans cells relies on autophagy-regulated lipid metabolism. J Cell Biol 224, e202403178 (2025).

10. Hara, T. et al. Suppression of basal autophagy in neural cells causes neurodegenerative disease in mice. Nature 441, 885–889 (2006).

11. Karasawa, K. et al. Vascular-Resident CD169-Positive Monocytes and Macrophages Control Neutrophil Accumulation in the Kidney with Ischemia-Reperfusion Injury. Journal of the American Society of Nephrology 26, 896 (2015).

12. Heib, T., Gross, C., Müller, M.-L., Stegner, D. & Pleines, I. Isolation of murine bone marrow by centrifugation or flushing for the analysis of hematopoietic cells – a comparative study. Platelets 32, 601–607 (2021).

13. Gautier, E. L. et al. Gene-expression profiles and transcriptional regulatory pathways that underlie the identity and diversity of mouse tissue macrophages. Nat Immunol 13, 1118–1128 (2012).

14. Chow, A. et al. CD169+ macrophages provide a niche promoting erythropoiesis under homeostasis and stress. Nat Med 19, 429–436 (2013).

15. Gupta, P. et al. Tissue-Resident CD169 + Macrophages Form a Crucial Front Line against Plasmodium Infection. Cell Reports 16, 1749–1761 (2016).

16. Kohyama, M. et al. Role for Spi-C in the development of red pulp macrophages and splenic iron homeostasis. Nature 457, 318–321 (2009).

17. Boada-Romero, E. et al. Membrane receptors cluster phosphatidylserine to activate LC3-associated phagocytosis. Nat Cell Biol 27, 1676–1687 (2025).

18. Heckmann, B. L. & Green, D. R. LC3-associated phagocytosis at a glance. Journal of Cell Science 132, jcs222984 (2019).

19. Stockwell, B. R. Ferroptosis turns 10: Emerging mechanisms, physiological functions, and therapeutic applications. Cell 185, 2401–2421 (2022).

20. Youssef, L. A. et al. Increased erythrophagocytosis induces ferroptosis in red pulp macrophages in a mouse model of transfusion. Blood 131, 2581–2593 (2018).

21. Cai, Z. et al. Loss of ATG7 in microglia impairs UPR, triggers ferroptosis, and weakens amyloid pathology control. J Exp Med 222, e20230173 (2025).

22. Ensinck, M. A., Brajovich, M. E. L., Borrás, S. E. G., Cotorruelo, C. M. & Biondi, C. S. Erythrocyte Senescent Markers by Flow Cytometry. OJBD 09, 47–59 (2019).

23. Vitaliti, A., Reggio, A. & Palma, A. Macrophages and autophagy: partners in crime. The FEBS Journal 292, 2957–2972 (2025).

24. Ganz, T. Macrophages and Iron Metabolism. Microbiology Spectrum 4, (2016).

