## Supplementary figures and images for "Autophagy deficiency in red pulp macrophages impairs their function and resistance to iron stress"

### Figure S1

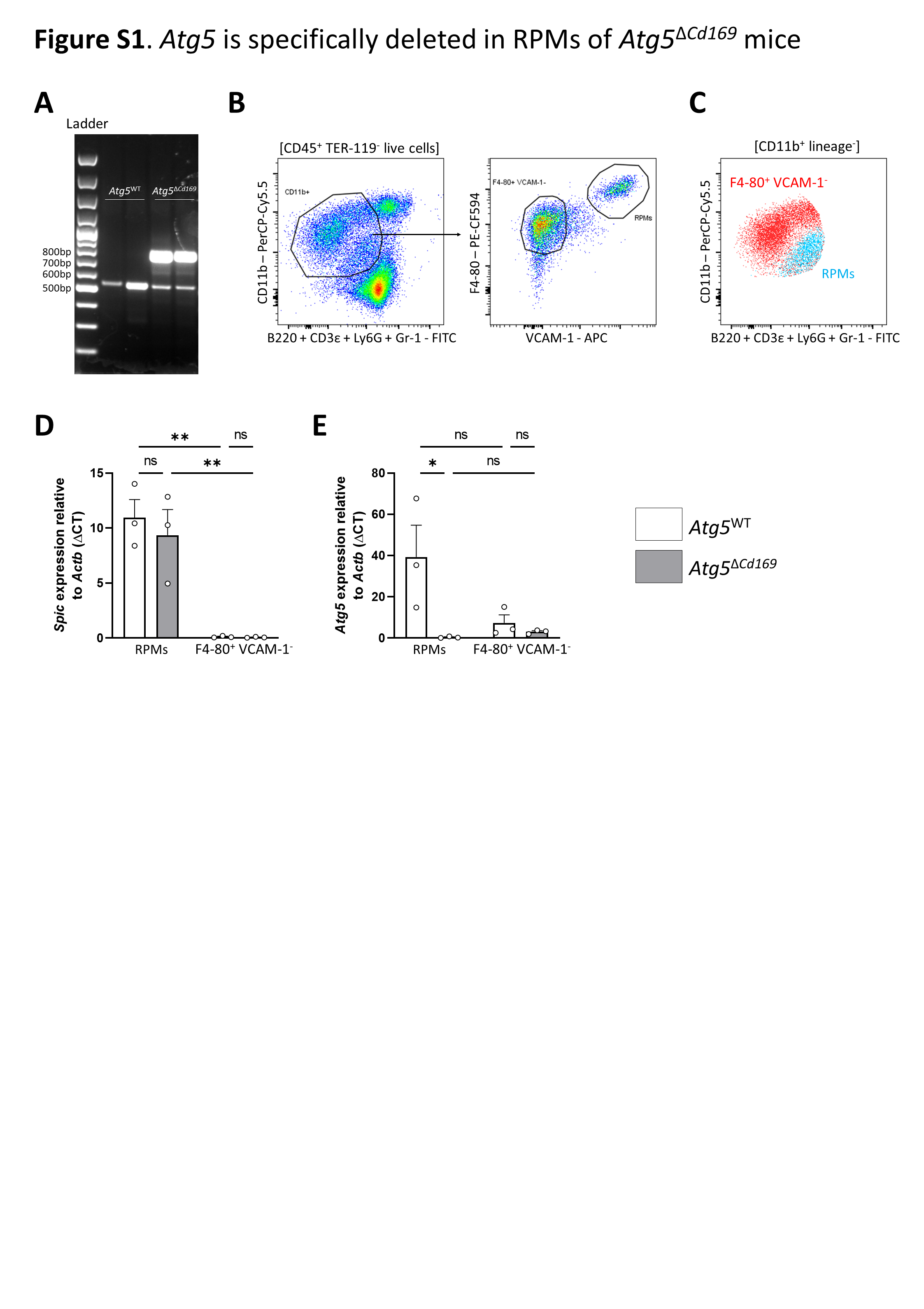

### Figure S2

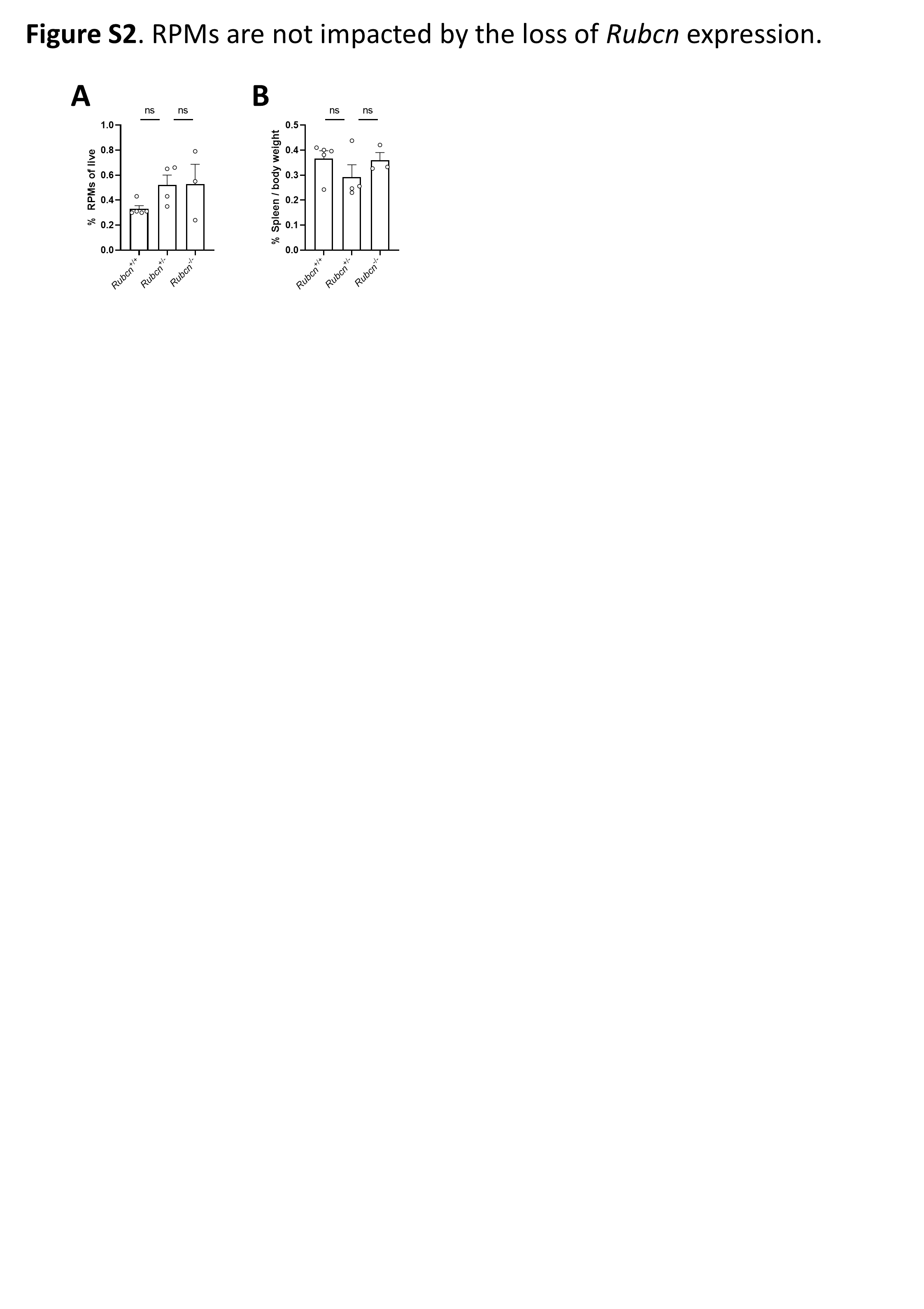

### Figure S3

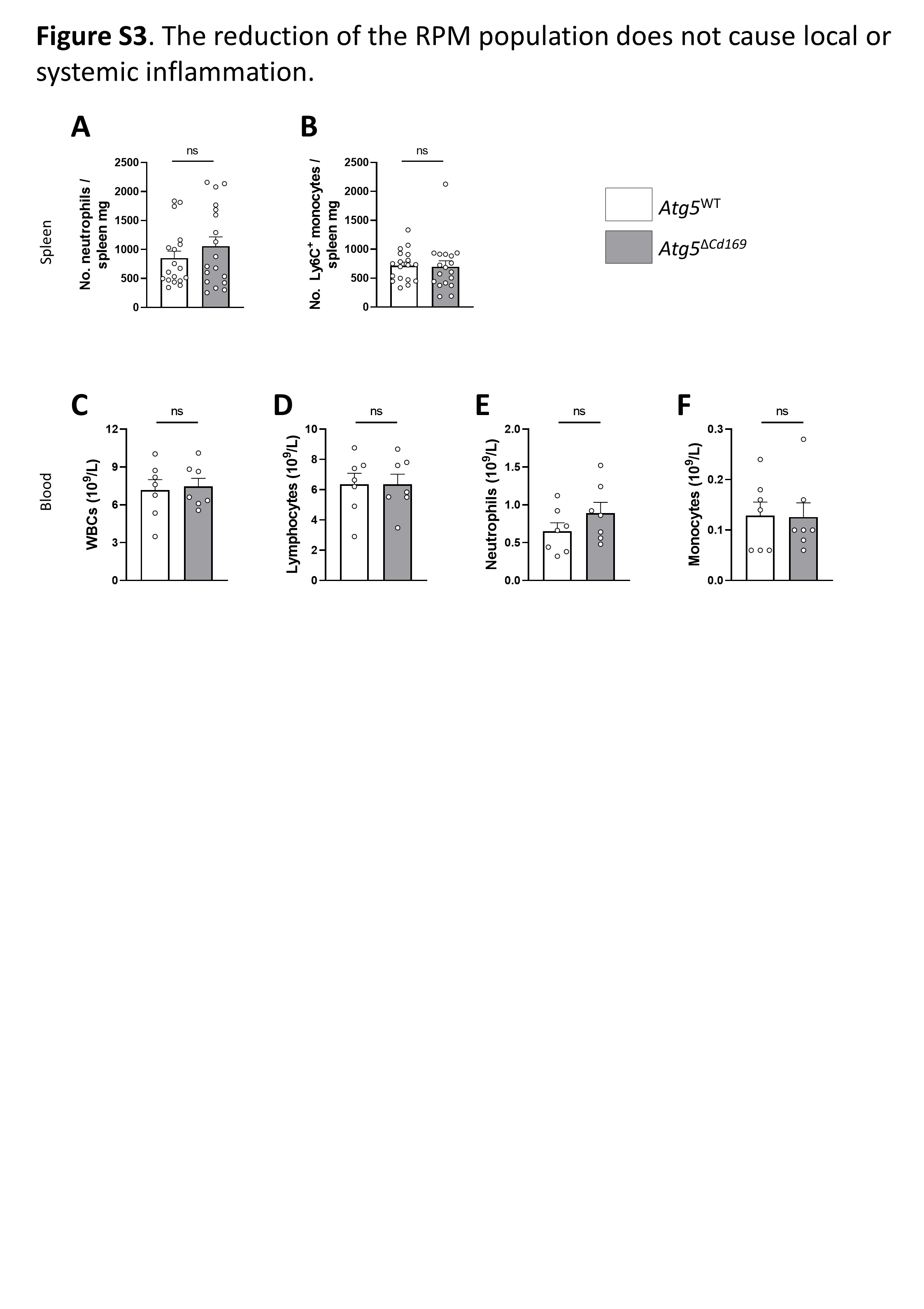

### Figure S4

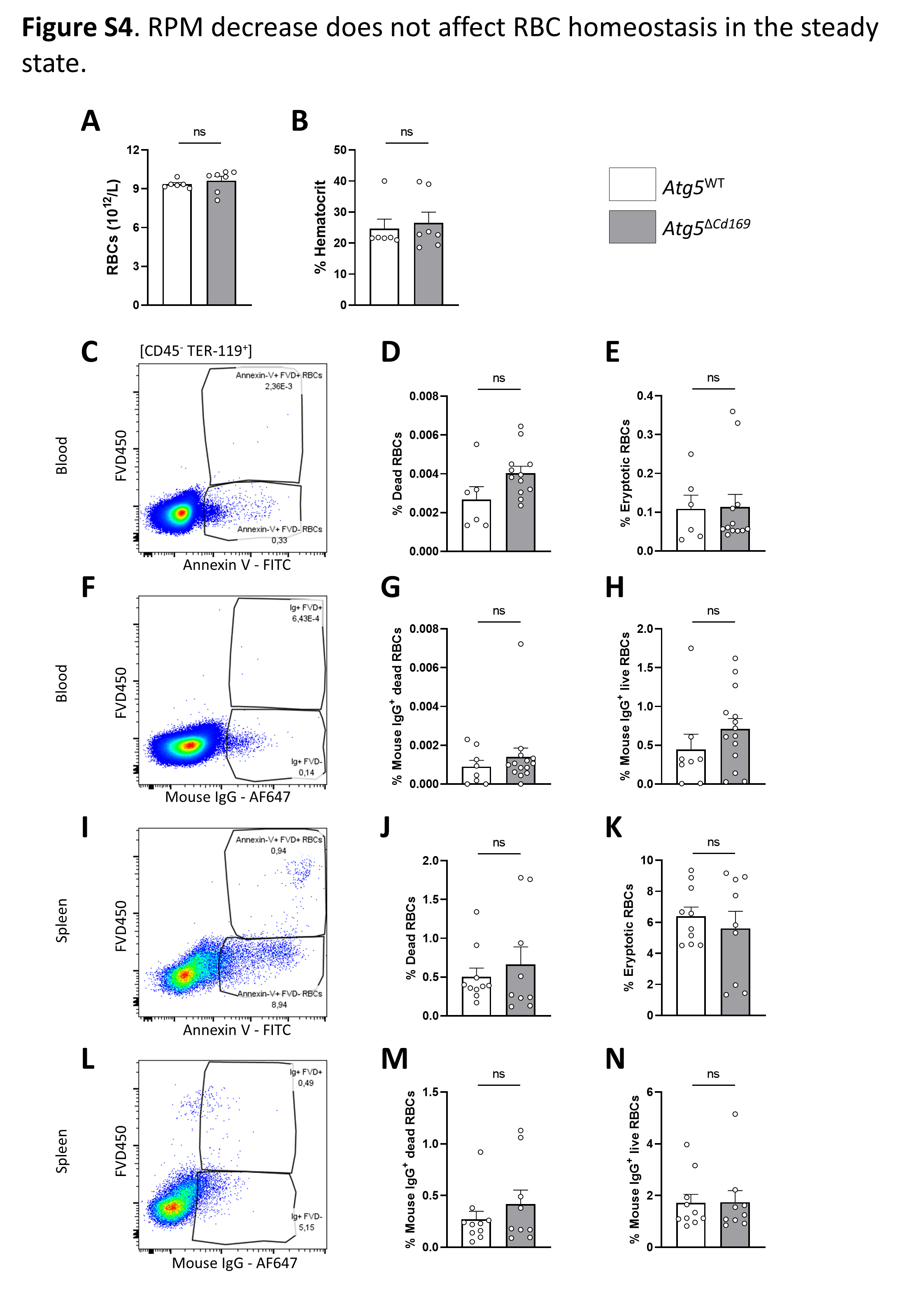

### Figure S5

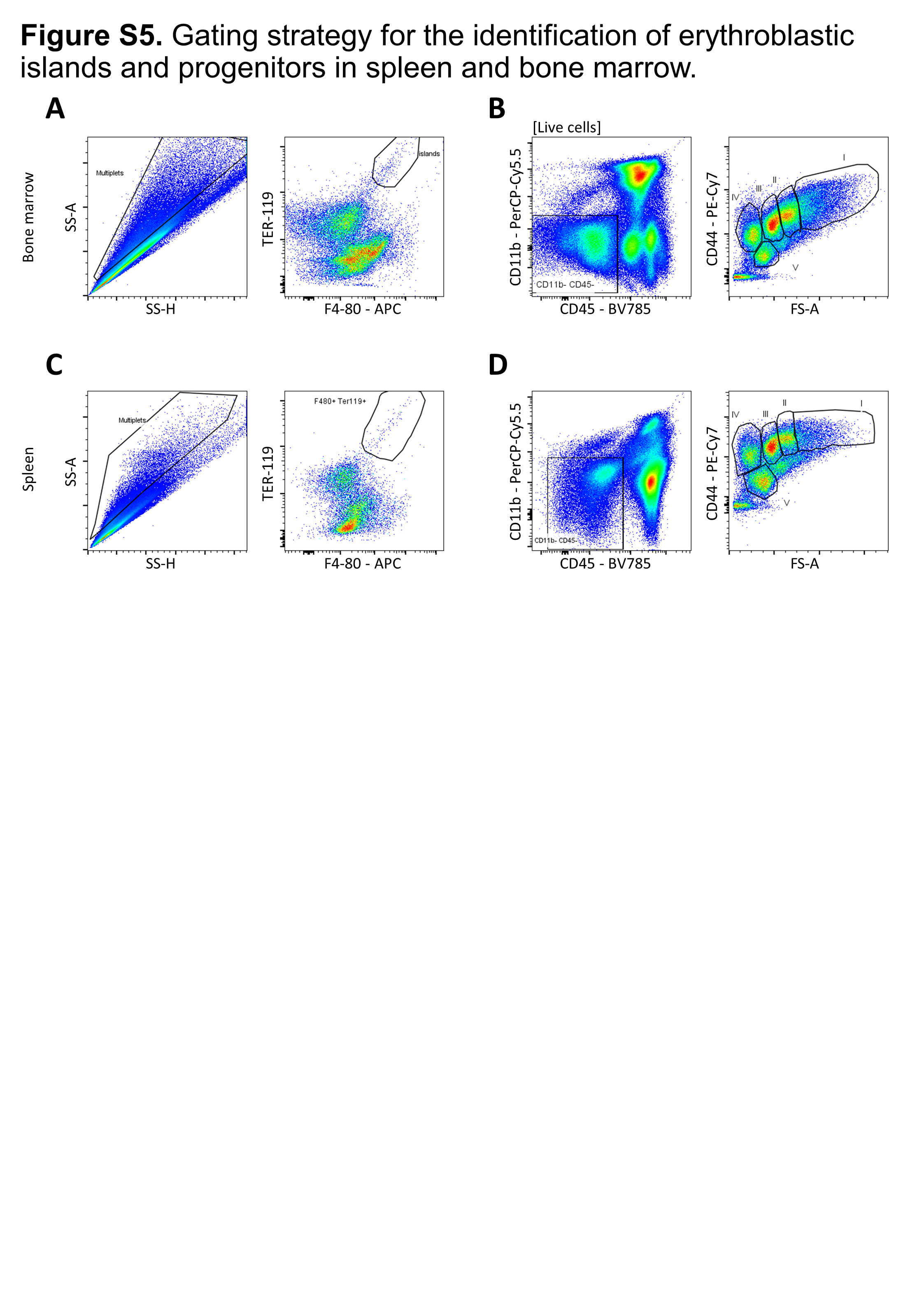
